# Global and Local Head Direction Coding in the Human Brain

**DOI:** 10.1101/2021.10.11.463872

**Authors:** J. P. Shine, T. Wolbers

## Abstract

Orientation-specific head direction (HD) cells increase their firing rate to indicate one’s facing direction in the environment. Rodent studies suggest HD cells in distinct areas of thalamus and retrosplenial cortex (RSC) code either for global (relative to the wider environment) or local (e.g., room-specific) reference frames. To investigate whether similar neuroanatomical dissociations exist in humans, we reanalysed functional magnetic resonance imaging data in which participants learned the orientation of unique images in separate local environments relative to distinct global landmarks (Shine, Valdés-Herrera, Hegarty, & Wolbers, 2016). The environment layout meant that we could establish two separate multivariate analysis models in which the HD on individual trials was coded relative either to global (North, South, East, West) or local (Front, Back, Right, Left) reference frames. Examining the data first in key regions of interest (ROI) for HD coding, we replicated our previous results and found that global HD was decodable in the thalamus and precuneus; the RSC, however, was sensitive only to local HD. Extending recent findings in both humans and rodents, V1 was sensitive to both HD reference frames. Additional small volume-corrected searchlight analyses supported the ROI results and indicated that the anatomical locus of the thalamic global HD coding was located in the medial thalamus, bordering the anterior thalamus, a region critical for global HD coding in rodents. Our findings elucidate further the putative neural basis of HD coding in humans, and suggest that distinct brain regions code for different frames of reference in HD.

**Significance statement:** Head direction (HD) cells provide a neural signal as to one’s orientation in the environment. HD can be coded relative to global or local (e.g., room-specific) reference frames, with studies suggesting that distinct areas of thalamus and retrosplenial cortex (RSC) code for this information. We reanalysed fMRI data where human participants associated images with global HDs before undergoing scanning. The design enabled us to examine both global and local HD coding. Supporting previous findings, global HD was decodable in thalamus, however the RSC coded only for local HD. We found evidence also for both reference frames in V1. These findings elucidate the putative neural basis of HD coding in humans, with distinct brain regions coding for different HD reference frames.

## Introduction

Determining one’s facing direction in the world is imperative for successful navigation. Head direction (HD) cells, found in brain regions including the thalamus and retrosplenial cortex (RSC), signal an animal’s facing direction via increased firing when the animal assumes the cell’s preferred orientation in the environment (Taube, 2007). With individual neurons signalling different HDs, the result is that of a neural compass providing essential input for other spatially-tuned neurons including place and grid cells (Harland et al., 2017; Winter, Clark, & Taube, 2015).

Theoretical models propose that HD cells form a ring attractor network (Knierim & Zhang, 2012). Cells proximal to one another on the ring signal similar facing directions and share excitatory interconnections, whereas distal cells, representing different directions, have inhibitory connections. This connectivity results in a bump of activity indicating the animal’s HD, which moves around the ring as a function of changes in orientation with respect to external landmarks and internally generated proprioceptive cues. This network structure has been observed in drosophila (Kim, Rouault, Druckmann, & Jayaraman, 2017) and other insects (Pisokas, Heinze, & Webb, 2020) but not more complex species, perhaps due to architectonically different arrangement of the ring attractor. Functional magnetic resonance image (fMRI) detects putative neural signals at the macroscopic level. As such, it is possible that HD cells distributed across cortex could contribute to a voxel-level signal, akin to the *in-vivo* study of grid cell signals in humans using fMRI (Stangl, Wolbers, & Shine, 2019).

Both global and local reference frames can be used to guide navigation. Behaviourally, there is evidence that while participants can integrate various route components into a single global reference frame (Mou, McNamara, & Zhang, 2013), local environment cues (e.g., the intrinsic geometry defined by the long axis of a room) also provide a reference frame for orientation (Mou & McNamara, 2002). At the neural level, in the RSC Jacob et al. (2017) found not only HD cells that tracked orientation relative to distal landmarks but also neurons tuned to the direction of local, room-specific environment cues. These findings accord with human multivariate fMRI results where activity patterns in the retrosplenial complex (a functionally-defined region extending beyond RSC) tracked information regarding local, but not global HD (Marchette, Vass, Ryan, & Epstein, 2014). More recently, Koch, Li, Polk, and Schuck (2020) found evidence for a multivariate global HD signal in RSC using a naturalistic navigation paradigm, supporting previous univariate results (Baumann & Mattingley, 2010; Shine et al., 2016). Similarly, the thalamus, a key HD region integrating body-based and visual information in rodents (Clark, Bassett, Wang, & Taube, 2010) and humans (Shine et al., 2016), is also hypothesised to represent distinct reference frames (Clark & Harvey, 2016) with anterior and lateral thalamus coding for global and local reference frames, respectively. These studies beg the question, however, of whether RSC and thalamus support concurrently distinct HD reference frames.

Despite its role in visual processing, converging evidence from rodent (Haggerty & Ji, 2015; Pakan, Currie, Fischer, & Rochefort, 2018; Saleem, Diamanti, Fournier, Harris, & Carandini, 2018), and human (Koch et al., 2020) studies suggest that area V1 codes also for HD. Although Koch et al. implemented analysis methods to decorrelate visual and HD information in a continuous navigation paradigm, a more stringent test of V1’s contribution to HD representations would separate the learning of HD, with its associated visual confounds, and the scanned test phase.

To address these questions, we used multivariate decoding to test for global and local HD signals across (sub)cortical structures previously implicated in HD coding. To achieve this aim, we re-examined data from Shine et al. (2016) where we observed univariate markers of HD coding in the thalamus, RSC, and precuneus. The study design provided us with the unique opportunity to explore global and local HD coding in the same task, as well as minimising visual confounds due to the separation of learning and the scanned test phases.

## Materials and methods

### Participants

Nine healthy volunteers (two female, seven male; age range 19-32), with normal or corrected-to-normal vision participated in the experiment at the UC Santa Barbara Brain Imaging Centre. The study received ethical approval from the local ethics committee.

### Procedure

For a detailed description of the paradigm, see Shine et al. (2016). Briefly, participants first explored a virtual environment (VE) outside of the MRI scanner whilst wearing a head mounted display (HMD). The environment comprised four galleries that were connected via a walkway, and four global landmarks (City, Bridge, and two mountain ranges) were rendered at infinity and provided cues to global HD. Each gallery contained a unique category of stimuli (Abstract art, Animals, Colour, Shapes), with the images on the walls inside the gallery aligned with the global landmarks in the environment. Participants were required to learn these individual images associated with the different global landmarks, which resulted in a unique quartet of images associated with each global HD (North, South, East, and West). The images inside each gallery were located either to the participant’s front, back, right or left upon entering the room. Facing direction in the environment was controlled via the participant physically rotating whereas translations were controlled via a button press.

After they had visited each gallery once, participants were tested on their knowledge of the environment via a judgement of relative direction task. Each of the 16 individual gallery images was presented separately and a text prompt asked the participants to point a joystick in the direction of a specific global landmark. Participants proceeded to the scanner phase of the experiment after they had passed the criterion (< 10 degree mean pointing error).

After successfully completing the learning, participants underwent scanning in which they were presented with the individual images from the galleries. Participants performed a one-back task regarding the associated global HD of each item, such that they pressed one button if the image was a repetition of the HD, and another button if HD was not repeated over consecutive trials. The 16 items were repeated twice per run, serving once as a non-repeat of HD and once as a repeat of HD. Importantly, gallery identity was not repeated (e.g., an image from the Abstract Art gallery never preceded itself). The scan session comprised eight runs, with images presented for five seconds and an inter-trial interval of 6-8 seconds. The design of the VE meant that it was possible to code trials according to different HD reference frames: 1) global head direction (GHD), in which images are coded according to their alignment with global environment landmarks, and 2) local head direction (LHD), in which images are coded according to their position relative to the participant when entering a gallery (Figure 1A). Importantly, repetition status (non-repeat and repeat) was balanced across the different conditions.

**Figure 1.**
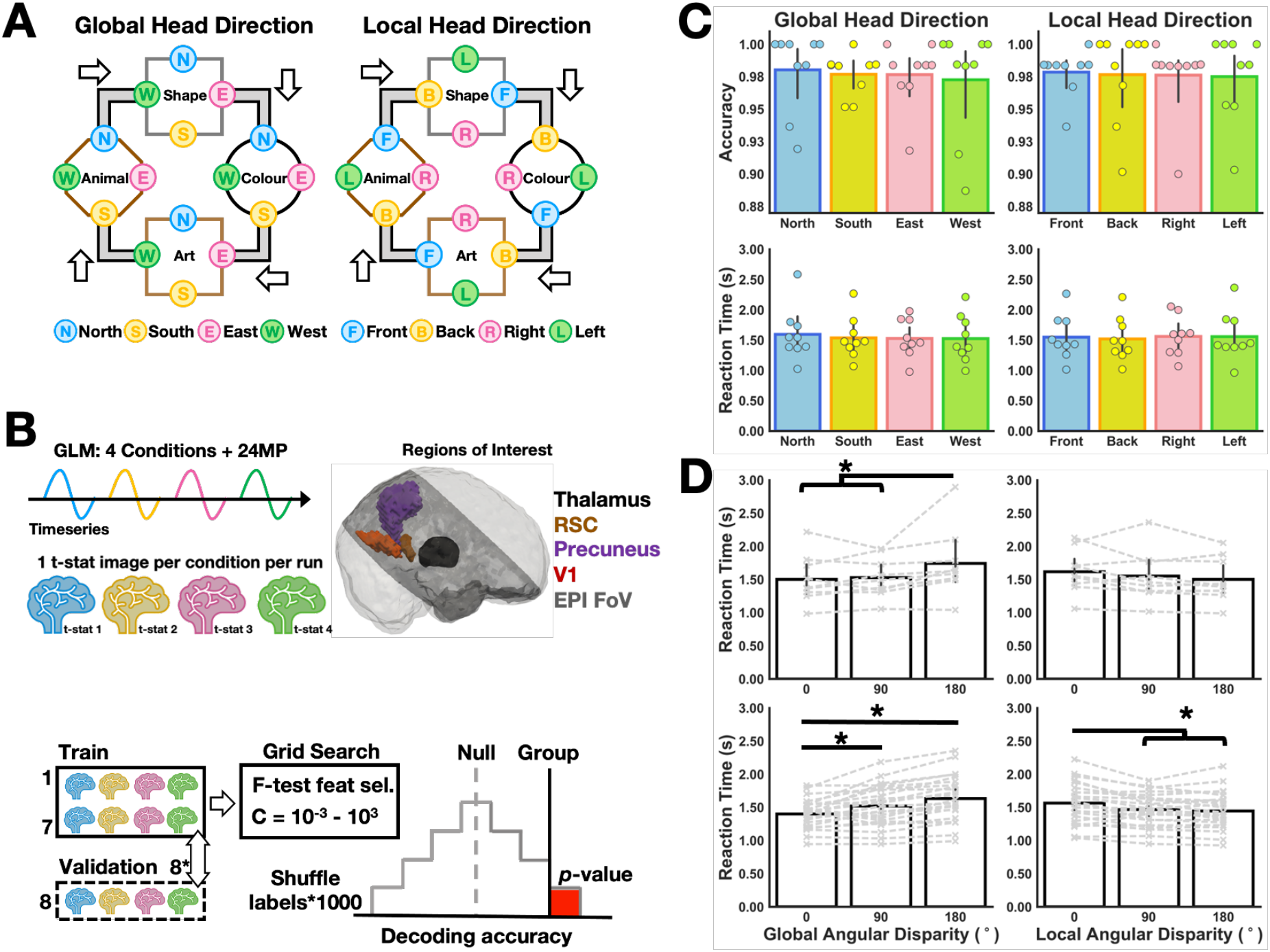
Overview of study design, preprocessing and behavioural performance. (A) The virtual environment consisted of four different galleries connected via walkways; the participant’s travel direction is indicated via the white arrows. Individual trials, comprising the four individual images inside each gallery, were coded for ‘Global Head Direction’ (i.e., the global reference frame defined by the external landmarks in the virtual environment; North, South, East, West), or ‘Local Head Direction’ (i.e., the local reference frame defined by the location of items as the participant entered the gallery; Front, Back, Right, Left), (B) Two separate general linear models (GLM) were created for the different reference frames. For each model, four regressors were used to model the conditions of interest, and the 24 motion parameters (MP) were added as nuisance regressors. This resulted in four t-stat images per run (32 images in total across eight runs), that were used for the decoding analysis in our key regions of interest for HD (i.e., thalamus, RSC, precuneus, V1). We used leave one-run-out nested cross-validation with a linear support vector classifier (SVC). To gain an unbiased estimate of decoding accuracy, the data were partitioned into train-test splits with one run left out to validate the accuracy of the model trained on the remaining data. In an inner loop, grid search was used to find the optimum combination of feature selection (using an F-test) and C-parameter for the SVC. This train-test split was repeated until all runs had served once as the validation dataset, and the mean score calculated per-participant, and for the group per-ROI. To determine the p-value associated with the group-level decoding accuracy, we created 1000 unique random shuffles of the condition labels and repeated the decoding analysis to generate a null distribution. The p-value was calculated as the number of times the decoding accuracy from the null distribution exceeded the observed group-level decoding accuracy, divided by the number of permutations. (C) Performance was matched across all conditions for each of the different models. (D) Upper panel: For the fMRI sample, grouping trials according to the angular disparity between consecutive trials revealed that reaction times were modulated in the GHD condition, such that differences of 180 degrees were significantly longer than 0 and 90 degrees. Lower panel: Repeating the paradigm in a larger behavioural sample (n=24) supported the results in the fMRI sample and showed a stepwise increase in RT according to the degree of GHD angular disparity. (* = p < 0.05); individual participant’s data displayed as a dashed grey line.

### Statistical analysis

Analysis of accuracy and reaction times (RTs) were conducted using R (version 3.6.3) with R-Studio (version 1.3.1056) and the afex package (version 0.28-0; Singmann, Bolker, Westfall, & Aust, 2018). Data were submitted to a repeated-measures ANOVA with four levels comprising either the GHD (North, South, East, West) or LHD (Front, Back, Right, Left). We tested also accuracy and RT as a function of the degree of angular disparity between subsequent trials in the run for both GHD and LHD. For example, if the first trial in the run was associated with GHD North, and the second trial GHD South, this second trial would comprise an angular disparity of 180 degrees. Relative to GHD North, subsequent GHD East or West trials were coded as 90 degrees. The same coding logic was applied to the LHD condition and the resulting values submitted to repeated-measures ANOVA with three levels (0, 90, 180 degrees) for GHD and LHD, respectively. For all analyses, where required, corrections for sphericity were made using Greenhouse-Geisser. Tukey’s post-hoc tests were used to interrogate significant main effects. Permutation-based analysis methods employed for the neuroimaging data are outlined below.

### MR imaging parameters

Imaging data were acquired using a 3T MRI scanner (Siemens Trio), with a standard receiver head coil. Twenty-five contiguous axial slices (3mm thickness), oriented to include key regions of interest (i.e., RSC, thalamus), were acquired using a T2*-weighted echo planar imaging (EPI) sequence (TR/TE = 1340/30ms; flip angle = 90 degrees; voxel size = 3mm isotropic, FoV = 192*192mm). A high resolution T1-weighted, whole-brain anatomical scan was acquired prior to the EPI data for each participant (TR/TE = 15/4.2ms; flip angle = 20 degrees; voxel size = 0.9mm isotropic, FoV = 240*240mm).

### fMRI data preprocessing and analysis

Data preprocessing was conducted using FSL v6.0 (FMRIB) (Jenkinson, Beckmann, Behrens, Woolrich, & Smith, 2012; Smith et al., 2004). The eight EPI runs were pre-processed as follows: The runs were concatenated, and a mean image created, and all eight runs were then realigned to the mean image, creating six motion parameters. The echoplanar imaging (EPI) data were skull-stripped, high-pass filtered, but left unsmoothed. For the region of interest (ROI) analyses, all analysis was performed in the participant’s native EPI space; for the whole-brain searchlight, functional data were non-linearly warped using FNIRT (Andersson, Jenkinson, & Smith, 2007) to the MNI-152 template.

To examine the two different spatial properties, three different general linear models (GLMs) were implemented, with each fMRI run modelled separately. In the GHD model, four regressors were created corresponding to whether the images in the galleries were aligned to the North, South, East, or West of the environment. In the LHD model, the four regressors reflected the arrangement of images as the participant entered the gallery (i.e., Front, Back, Right or Left). All regressors were convolved with a double-gamma HRF. In addition to the four regressors described for each model, each GLM included the six motion parameters, their derivatives, and the squares of these values (24 motion regressors in total) as nuisance regressors. For each run the resulting t-stat images associated with the four conditions in each separate model were used for the decoding analysis. A suite of Python (version 3.6.7) packages including pandas (version 0.20.3; McKinney, 2010) and NumPy (version 1.14.2; Harris et al., 2020) were used to analyse the data. NiBabel (version 2.1.0; Brett et al., 2019) was used to load the t-stat images, and the decoding analysis carried out using Scikit-learn (version 0.18.1; Pedregosa et al., 2012). Decoding results were plotted using Seaborn (version 0.7.1; Waskom et al., 2016).

In line with the best practices of decoding (Varoquaux et al., 2017), we used a nested-cross validation procedure with a linear Support Vector Classifier (SVC) where we determined the best performing decoding parameters via grid search to obtain an unbiased estimate of decoding accuracy. The decoding analysis pipeline comprised Z-scoring the data across voxels, univariate feature-selection via an F-test, and multivariate decoding using a Linear SVC. For the F-test, we selected either the voxels with the 20^th^ percentile highest F-scores, or all voxels (i.e., the 100^th^ percentile) in the region of interest (ROI). For the linear SVC, the C-parameter ranged from 10^-3^ to 10^3^ in steps of 10^1^. To determine an unbiased estimate of decoding accuracy, we used nested cross-validation where one of the eight runs of data was partitioned as the validation-set and not used for tuning of the model’s hyper-parameters (i.e., feature-selection F-test percentile and SVC C-value). For the remaining seven runs, the data were partitioned into seven train-test splits, and all combinations of the F-test and C-parameter values crossed to determine the combination of best performing parameters. These values were then fit on the seven runs and used to predict class label in the unseen validation-set. This procedure was repeated eight times so that every run served as a validation set. The scores of these eight runs were then averaged to generate a participant-specific decoding accuracy and averaged across participants to determine the group-level decoding accuracy.

To determine the p-value associated with the group-level decoding accuracy, we generated 1000 shuffled orders (without repetition) of the condition labels that respected the frequency of target labels per run. We repeated the analysis pipeline with each of the shuffled orders, resulting in a null distribution of 1000 group-level decoding accuracies. The *p*-value was calculated as the number of times the null group-level decoding accuracies equalled or exceeded the observed group-level decoding accuracy, divided by the number of permutations; one was added to both the numerator and denominator of this calculation to ensure that the p-value never equalled 0 (an overview of the analysis method is shown in Figure 1B).

The decoding analyses were carried out in regions of interest (ROIs) that have been shown to be involved in HD coding in humans. Specifically, we examined decoding in the thalamus segmented using FSL’s FIRST, the RSC derived from Freesurfer (Shine et al., 2016), and probabilistic masks of V1 and precuneus from the Harvard-Oxford subcortical atlas; probabilistic masks were thresholded at 50% likelihood of the target ROI. Anatomical masks of thalamus and RSC were warped to the subject’s native EPI-space via the inverse of the EPI-to-T1 Affine matrix, and the probabilistic masks were warped from MNI-space using the inverse of the EPI-MNI matrix.

We implemented also a whole brain searchlight to examine brain regions outside of our target ROIs where we could reliably decode the different spatial properties. The searchlight procedure followed that detailed by Stelzer, Chen, and Turner (2013). Specifically, we used a searchlight radius of 6mm (i.e., two voxels, which has been shown to perform best in searchlight analyses; Kriegeskorte, Goebel, & Bandettini, 2006) and ran the same decoding analysis pipeline detailed above, with the exception that the SVC C-parameter was fixed at one to reduce the computational burden. First, for each participant we ran the whole brain searchlight with the correct condition labels where the decoding accuracy was assigned to the central voxel of the searchlight sphere. This provided a whole brain decoding accuracy map. The individual participant accuracy maps were then moved to standard MNI space and averaged across participants to obtain a whole brain, group-level decoding accuracy map. To generate the null distribution with which to compare the group-level decoding accuracy map, we first repeated the searchlight analysis with 100 permutations of the condition labels for each participant. The 100 permuted accuracy maps were then also moved to MNI space. We created a mask that contained voxels in which all participants’ data overlapped and used this to mask the searchlight results. We then randomly selected one of the 100 permutations (with replacement) per participant and averaged these values to generate a group-level permuted accuracy map. This random selection was repeated 100,000 times resulting in 100,000 permuted accuracy maps that formed the basis for our null distribution. For each voxel we had 100,000 accuracy values comprising the voxel-wise null distribution, and we used a threshold of p < 0.01 to determine an accuracy threshold that had a low probability of occurring by chance. These voxel-wise threshold accuracy values were then applied to both the group level decoding accuracy map as well as the 100,000 permuted group accuracy maps. Voxels surviving the threshold were binarized and cluster statistics performed on the group level decoding accuracy map, as well as each of the permuted group accuracy maps, using FSL’s cluster; a cluster comprised two or more voxels that shared a face (i.e., connectivity = 6). We recorded all resulting clusters and their frequency from the permuted group accuracy maps, which allowed us to estimate the probability of observing by chance a cluster of a given size in the group level decoding accuracy map (i.e., number of clusters of a given size larger than our observed cluster divided by the total number of clusters in the permuted accuracy maps). To correct for multiple comparisons, for all the clusters of the group level decoding accuracy map, we applied a step-down FDR correction (Benjamini & Liu, 1999) as implemented in R package Mutoss (version 0.1-12). Whole brain searchlight results were rendered on the MNI-152 template using MRIcroGL (version 1.2.20200331).

## Results

### Behavioural

Participants were trained to discriminate GHD to a high level of accuracy outside of the scanner. Reflecting this training, participants retained this high level of performance on the 1-back task during the scanner task (Figure 1). Importantly, for decoding, accuracy (F(1.74, 13.88) = 0.24, p = 0.76, η^2^ = 0.009) and reaction times (F(1.85, 14.76) = 0.56, p = 0.57, η^2^ = 0.007) were matched across the four different GHDs (Figure 1C). Coding the trials according to LHD did not affect the pattern of data in terms of accuracy and reaction times (all Fs < 0.35, ps > 0.69, η^2^ < 0.003).

According to the architecture of ring attractor networks (Knierim & Zhang, 2012; Zhang, 1996), HD cells coding for similar orientations predominantly share excitatory connections whereas cells coding for very disparate orientations share more inhibitory connections. Ring attractor dynamics would therefore predict that reaction times should vary as a function of the degree of angular disparity between consecutive trials. For example, if a given trial was associated with GHD north (0 degrees) and the next trial was associated with GHD south (180 degrees), this angular disparity of 180 degrees should be harder to overcome than a disparity of 90 degrees. We performed this analysis both for GHD and LHD (Figure 1D). For GHD, we observed a main effect of angular disparity (F(1.41, 11.31) = 6.32, p = 0.02, η^2^ = 0.075), which was driven by reduced reaction times for 0 versus 180 (t(16) = −3.25, p = 0.013), and 90 versus 180 degrees (t(16) = −2.87, p = 0.028); angular disparities 0 and 90 degrees did not differ (t(16) = −0.38, p = 0.924). LHD reaction times were not influenced by the angular disparity between subsequent trials (F(1.29, 10.35) = 2.51, p = 0.14, η^2^ = 0.02).

To test the veracity of our behavioural results, we repeated the paradigm in an additional sample of 24 participants (14 female, 10 male; mean age: 24.2, age range: 20–31). The paradigm was identical except that the intertrial interval was reduced to two seconds. Examining first the GHD, we found a significant main effect of angular disparity (F(1.33, 30.5) = 24.246, p < 0.001, η^2^ = 0.098), with a step-wise increase in RT according to increasing angular disparity (Figure 1D). Specifically, 0 degree changes in GHD were significantly faster than 90 (t(46) = −3.53, p = 0.003) and 180 degree changes (t(46) = −6.96, p < 0.0001); 90 degree changes in GHD were also significantly quicker than 180 degrees (t(46) = −3.43, p = 0.004). The same analysis for LHD revealed a main effect of angular disparity F(1.59, 36.54) =13.58, p = 0.0001, η^2^ = 0.034). This main effect was driven by significantly longer responses for 0 versus 90 and 180 degree changes in LHD (t(46) = 4.01, p = 0.0006 and t(46) = 4.89, p < 0.0001, respectively); 90 and 180 degree changes in LHD did not differ (t(46) = 0.88, p = 0.66).

### Decoding in ROIs

Our multivariate analyses first focussed upon regions of the brain previously shown to be involved in HD coding, namely the thalamus, RSC, precuneus, and V1. For each ROI, we examined the decoding accuracy of the two different reference frames (GHD and LHD) and calculated the corresponding p-value derived from a null distribution generated via 1000 permutations of the condition labels. For GHD (Figure 2), we could decode different directions relative to the distal landmarks with above chance accuracy in bilateral thalamus (p = 0.007), precuneus (p = 0.048) and V1 (p = 0.001). In contrast to our previous univariate results (Shine et al., 2016), there was no evidence of multivariate signal for GHD in RSC (p = 0.419). Next, we examined the coding of gallery-specific LHD in the same ROIs (Figure 3). Unlike GHD, it was possible to decode LHD in RSC (p = 0.049). In line with the results for GHD, V1 contained information also regarding LHD (p = 0.002). It was not possible, however, to decode LHD in thalamus (p = 0.266) or precuneus (p = 0.237).

**Figure 2.**
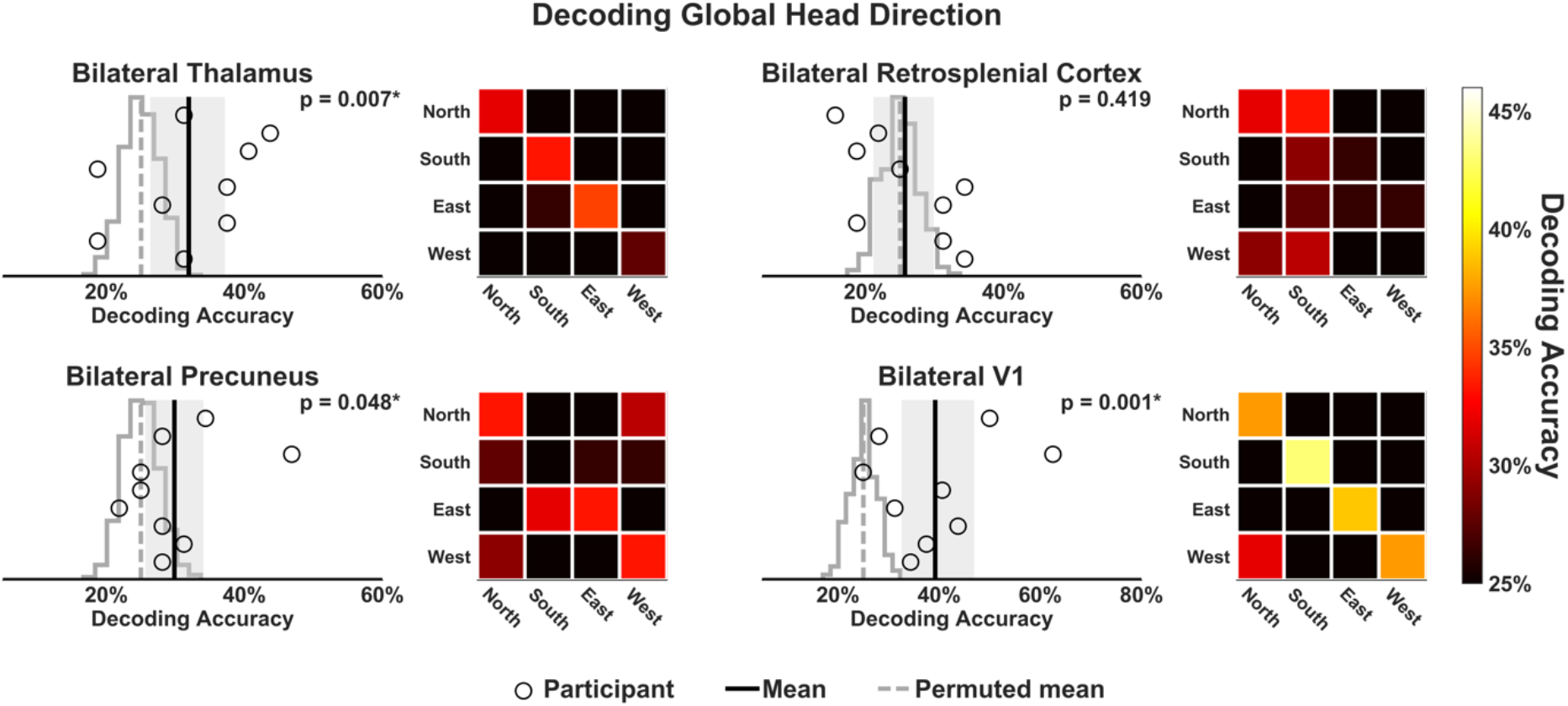
Decoding Global Head Direction in key HD ROIs. The linear SVC revealed significantly above chance decoding accuracy of GHD in bilateral thalamus (top left), precuneus (bottom left) and V1 (bottom right). Individual subject’s data are displayed as black circles, with the mean group decoding accuracy denoted by the solid black vertical line. The grey shading indicates the 95% confidence interval around the mean group accuracy, calculated using bias-corrected and accelerated bootstrap with 1000 samples with replacement of the subjects’ decoding scores. The confusion matrices, located to the right of each plot, show the mean group decoding accuracy separated according to the different HDs.

**Figure 3.**
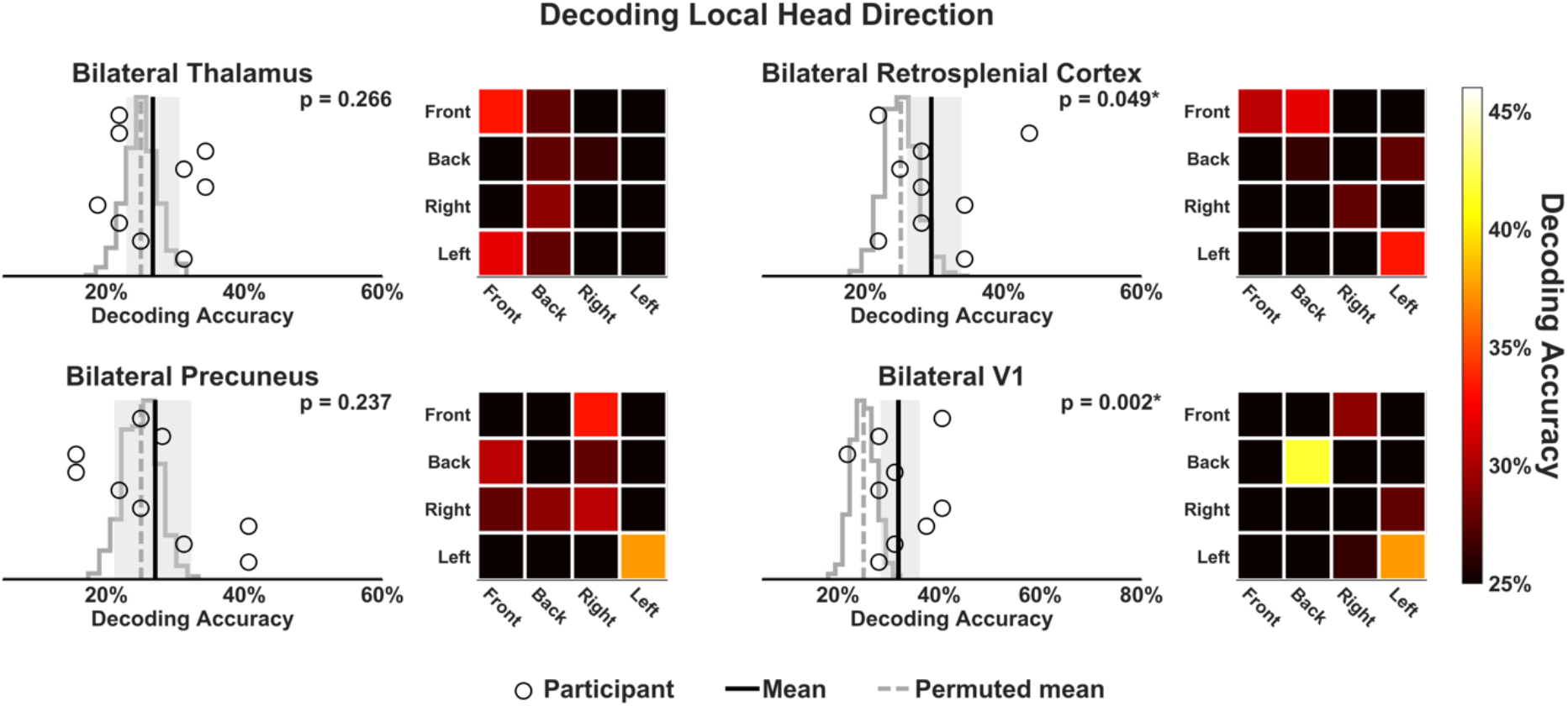
Decoding Local Head Direction in key HD ROIs. The linear SVC revealed significantly above chance decoding accuracy of LHD in bilateral RSC only (top right). Individual subject’s data are displayed as black circles, with the mean group decoding accuracy denoted by the solid black vertical line. The grey shading indicates the 95% confidence interval around the mean group accuracy, calculated using bias-corrected and accelerated bootstrap with 1000 samples with replacement of the subjects’ decoding scores. The confusion matrices, located to the right of each plot, show the mean group decoding accuracy separated according to the different HDs.

### Searchlight analysis

To examine whether regions of cortex outside of our ROIs code for different HD reference frames, we conducted whole brain decoding using a searchlight sphere with a radius of 6mm and implemented the same decoding scheme. For GHD, four clusters survived FDR correction at the whole brain level. Consistent with the ROI analyses, one cluster was located in the precuneus (Figure 4), while another was located inferior to the precuneus and posterior to the RSC. The remaining two clusters were located in the lateral occipital cortex and occipital pole (Figure 5). The same analysis conducted for LHD did not yield any significant clusters.

**Figure 4.**
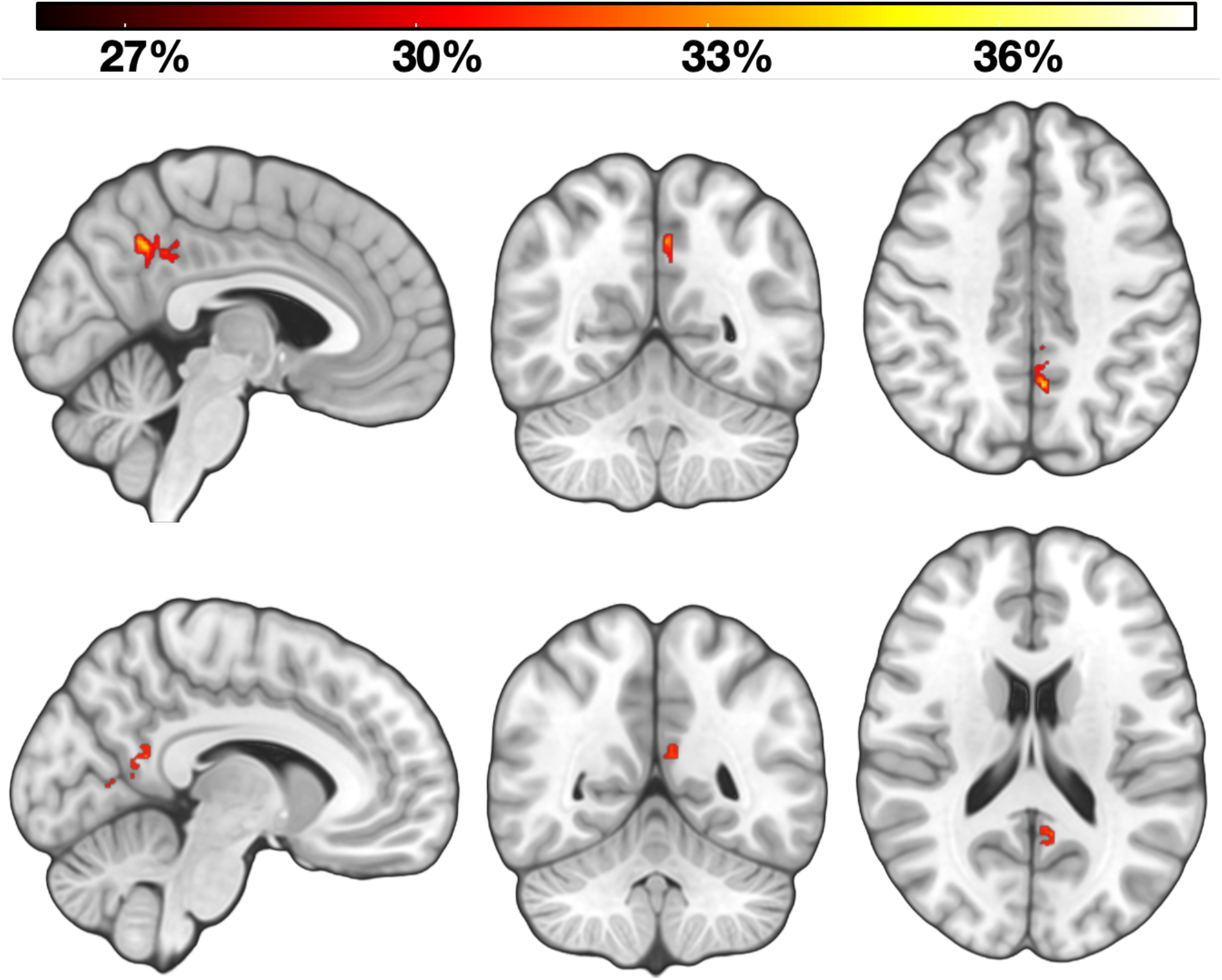
Whole brain searchlight decoding of Global Head Direction in precuneus/posterior medial cortex. Using a whole brain searchlight we found two clusters of above chance decoding accuracy in the posterior medial cortex. Upper panel: One cluster was located in the precuneus/posterior cingulate (−4, −56, 42; 80 voxels; peak decoding accuracy = 36.8%; FDR-adjusted *p*-value = 0.007). Lower panel: the other cluster was located posterior to the retrosplenial cortex (−8, − 52, 18; 63 voxels; peak decoding accuracy = 35.4%; FDR-adjusted *p*-value = 0.02). The heatmaps reflect percent decoding accuracy.

**Figure 5.**
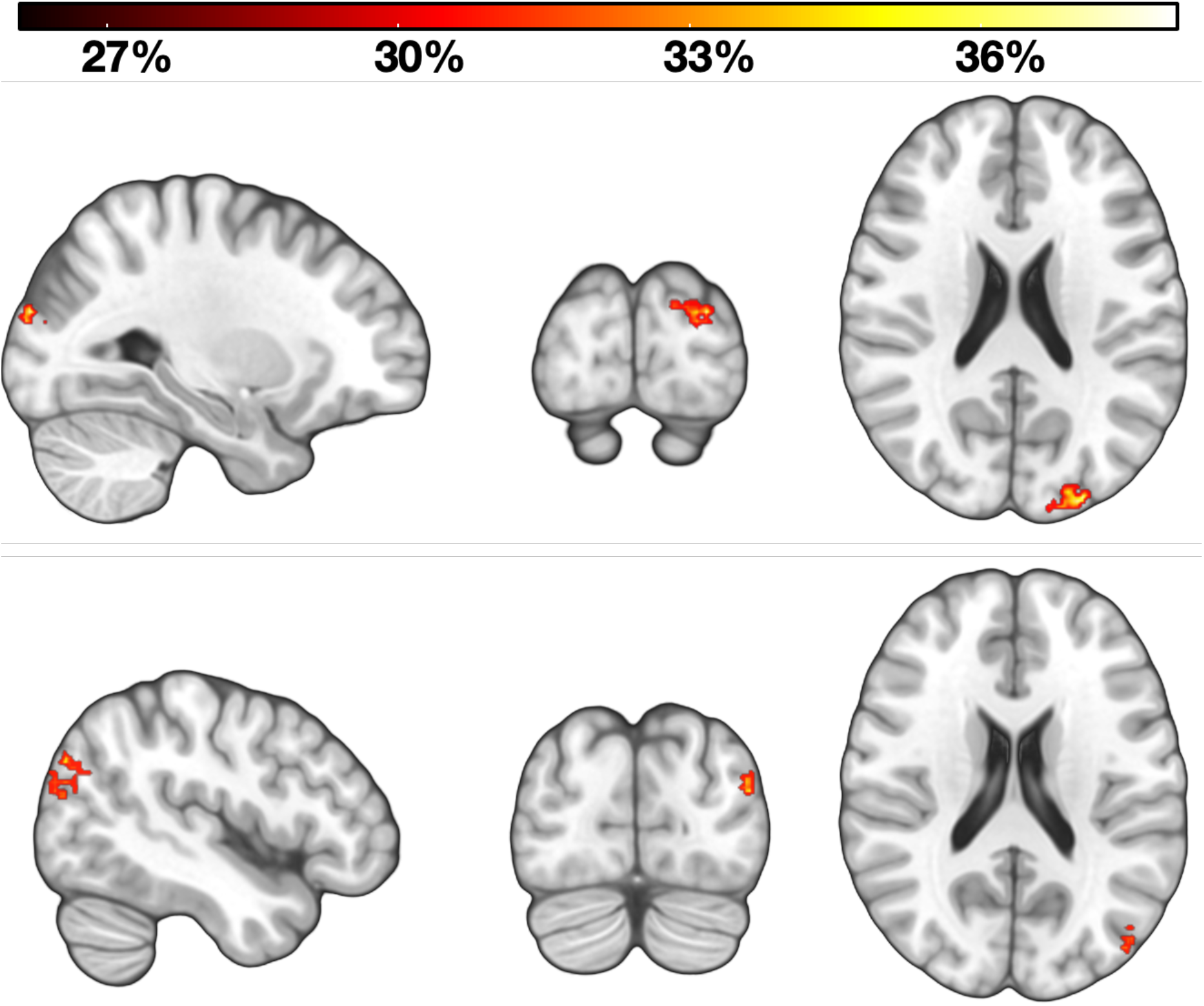
Whole brain searchlight decoding of Global Head Direction. Two additional clusters of above-chance decoding accuracy were identified in the whole brain searchlight analysis. Upper panel: Occipital pole/Lateral occipital cortex (−28, −92, 22; 146 voxels; peak decoding accuracy = 38.9%; FDR-adjusted p-value = 0.0002). Lower panel: Lateral occipital cortex (−46, −82, 20; 114 voxels; peak decoding accuracy = 37.2%; FDR-adjusted p-value = 0.0008). The heatmap reflects percent decoding accuracy.

To provide greater insight into the anatomical location of the GHD coding effects located in the thalamus in the ROI analysis, we applied a small volume correction and re-ran the searchlight in the bilateral thalamus mask. Using the same 6mm searchlight radius, we found a significant cluster of above chance decoding accuracy located in left medial thalamus (Figure 6). To ensure that the result was not an artifact of the searchlight radius (Etzel, Zacks, & Braver, 2013), we repeated the analysis in the thalamus with different searchlight radii to examine the consistency of the results. At 3mm, the decoding accuracy was not significant (p = 0.07), but with searchlight radii larger than the 6mm used initially (i.e., at 9mm and 12mm) we again identified significant clusters in the left medial thalamus, bordering the anterior thalamus (Figure 6). Despite the absence of any significant effects for LHD in the whole brain searchlight, for completeness, we conducted the same analysis for LHD in thalamus. This analysis, however, did not reveal any significant clusters regardless of the searchlight radius.

**Figure 6.**
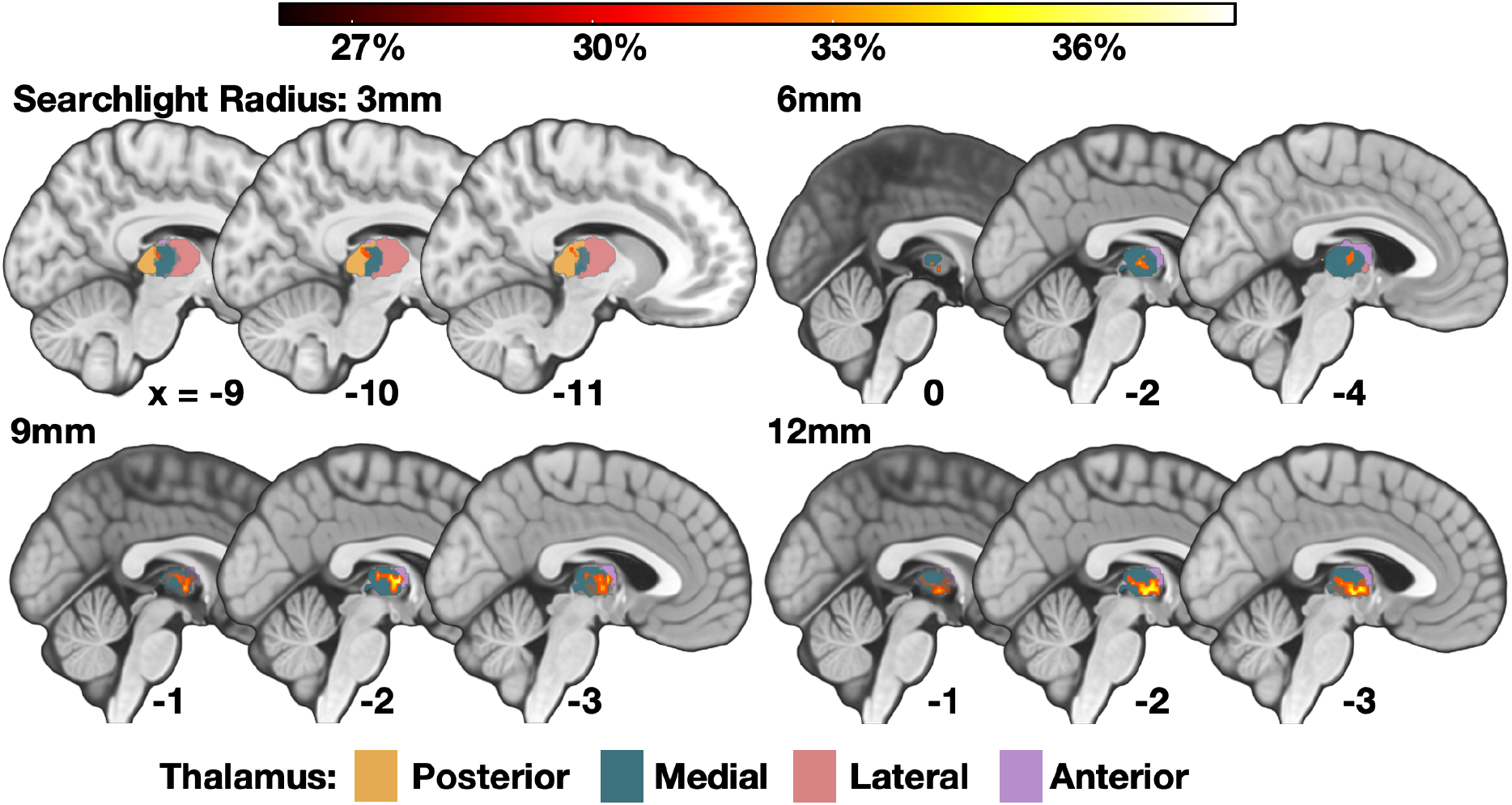
Small-volume-correction decoding of Global Head Direction in bilateral thalamus using different searchlight radii. Using a small volume correction, we found a significant cluster of above chance decoding accuracy located in the left thalamus using a 6mm searchlight radius, consistent with the whole brain methods (−2, −6, 6; 18 voxels; peak decoding accuracy = 34.7%; FDR-adjusted *p*-value = 0.044). To examine the reliability of this effect, we conducted the same analysis using additional searchlight radii (3, 9, and 12mm). For the 3mm searchlight, we could not reliably decode GHD (0, −10, −2; 7 voxels; peak decoding accuracy = 33.7%; FDR-adjusted *p*-value = 0.07). For both the larger 9 and 12mm radii, however, there was evidence of GHD information located in medial thalamus, bordering the anterior thalamus (9mm: −2, −6, 6; 69 voxels; peak decoding accuracy = 36.8%; FDR-adjusted *p*-value = 0.016; 12mm: −2, −12, −2; 131 voxels; peak decoding accuracy = 36.8%; FDR-adjusted *p*-value = 0.018). Probabilistic maps of thalamic subregions created from (Krauth et al., 2010). Values beneath each sagittal slice indicate *x* coordinate. The heatmaps reflect percent decoding accuracy.

## Discussion

We explored the contribution of key ROIs involved in HD to the decoding of different spatial reference frames by performing a reanalysis of the data from Shine et al. (2016). Extending previous findings, we found that the human thalamus contains a multivariate signal of GHD but not LHD. Consistent with univariate results, this GHD information was present also in the precuneus, and, in line with recent findings (Koch et al., 2020), V1. In RSC, however, it was possible to decode only the LHD. As well as GHD, LHD information was also evident in V1.

In line with our previous findings (Shine et al., 2016), our results suggest that the thalamus contains a GHD signal. Our searchlight analyses indicate that this effect located in the left medial thalamus, bordering the anterior thalamus - a region comprising the anterior dorsal thalamic nuclei that contain HD cells in rodents. That the results were consistent at several different, but not all, searchlight radii suggests that the GHD results were not simply an artifact of the searchlight size. These data may reflect the coarse organisation of a ring attractor network in the thalamus, or at least a signal that can be extracted at the macroscopic level. Future rodent studies that sample a large number of neurons from the HD circuit will help elucidate possible patterns of network organisation of the HD system, although it will likely differ substantially across species (Pisokas et al., 2020). We observed also a pattern of behavioural data consistent with the predictions one might make based upon the ring attractor architecture, with greater angular disparity between consecutive trials’ GHDs resulting in longer RTs, which we replicated in a larger behavioural sample. It is possible that such readouts might provide greater insight into ring attractor structure and/or integrity in humans.

In contrast to GHD, there was no evidence that the thalamus codes for LHD as previously hypothesised (Clark & Harvey, 2016). This was true even when using a whole brain searchlight analysis. Although we must be cautious when interpreting the absence of an effect, one possible explanation as to why coding for LHD was not apparent is that participants were trained in our task to focus upon the global landmarks for HD. Despite these task demands it should be noted, however, that we did see decoding of LHD in other brain regions (i.e., RSC/V1). It could be the case that HD representations in different regions are modulated to a greater or lesser degree by attentional demands, and that representations in the thalamus change as a function of the reference frame required to solve the current navigational problem. Future studies that manipulate attention to different reference frames will be required to examine this question.

Unlike the univariate results observed in Shine et al. (2016), there was no evidence of a GHD signal in RSC. Consistent with Marchette et al. (2014), however, we could decode LHD. Our findings are at odds with recent data from Koch et al. (2020), in which they decoded walking direction in RSC. In their study, the environment comprised a single open arena and did not require the integration of different LHDs in service of a GHD. It could be argued, therefore, that this RSC effect represents the coding of a single local reference frame. The apparent inconsistencies between the results of the univariate analysis in Shine et al. (2016) and those reported here could reflect the structure of the RSC and the arrangement of the HD cells in the granular and dysgranular cortex. In Jacob et al. (2017), bidirectional HD cells were detected almost exclusively in the dysgranular RSC, whereas traditional HD cells, which were sensitive to the global cues, were found in both granular and dysgranular RSC. The current fMRI resolution means that it is not possible to discriminate substructures of RSC, however an intriguing hypothesis from the current data is that there is greater structure in the arrangement of orientations for cells coding for LHD relative to GHD. This coarse organisation would mean that, at the voxel-level, a distinct signal would be evident for each LHD, resulting in above-chance decoding (Drucker & Aguirre, 2009; Epstein & Morgan, 2012). Combined with the univariate results of Shine et al. (2016), the RSC contains information regarding both HD reference frames, and mirrors the rodent data in which neurons in the RSC have been shown to encode diverse spatial properties, such as left versus right hand turns in a maze, as well as the animal’s position on a track in both ego- and allocentric coordinates (Alexander & Nitz, 2015; Nitz, 2012).

The successful decoding of GHD and LHD in V1, despite the stimuli being balanced in terms of stimulus category, adds to the growing evidence of the contribution of primary visual regions to spatial processing in rodents (e.g., Pakan et al., 2018; Saleem et al., 2018) and humans (Koch et al., 2020). Importantly, however, our learning and test phases were separate, meaning that visual information (e.g., a visible global landmark in the arena) was less likely to play a role in our findings. One possible explanation is that the decoding of GHD in V1 reflects representations of the associated global landmark (e.g., bridge, mountain). This explanation, however, does not accommodate the findings for LHD, in which both stimulus category and global landmark were balanced. The decoding of GHD in V1 could reflect reinstatement of visual information associated with a given facing direction, driven by, for example, the multimodal visuo-vestibular representation of GHD in the thalamus. This explanation is perhaps unlikely, however, given that spatial signals in the rodent V1 persist even in the absence of modulation of thalamic input from the lateral geniculate nucleus (Diamanti et al., 2021). Future studies that acquire high-resolution temporal information may help to shed light on the origin of this HD signal in V1, but it seems apparent from several studies that V1 plays a more prominent role in spatial coding than previously anticipated.

In addition to the ROI data, we observed at the whole brain level via searchlight a GHD signal in the precuneus, which supports the notion that this parietal region contains world-centred coordinates (Snyder, Grieve, Brotchie, & Andersen, 1998; Wilber, Clark, Forster, Tatsuno, & McNaughton, 2014). Furthermore, we found a significant cluster in the posterior medial cortex, posterior to the RSC, in a region analogous to the retrosplenial complex where coding of HD has been reported previously (Baumann & Mattingley, 2010; Vass & Epstein, 2013). The relative frequency of the effects in these posterior medial regions, not limited to anatomical RSC, highlights the importance of these regions in spatial coding. In terms of clinical applications, recent studies have shown that the posterior medial cortex is an early target for amyloid deposition (Maass et al., 2019) and shows altered function in young adults at risk of Alzheimer’s Disease (Shine, Hodgetts, Postans, Lawrence, & Graham, 2015). That these regions are impacted in the early stages of Alzheimer’s and appear involved in GHD coding may help to explain spatial disorientation observed in the disease (Lester, Moffat, Wiener, Barnes, & Wolbers, 2017).

There are limitations to the conclusions that can be made on the basis of these data. First, this study comprised the reanalysis of data with a small sample size. Despite using non-parametric analysis methods to lessen the impact of the sample size, it would be important to see the results replicated in a larger sample. The findings, however, appear consistent across univariate and multivariate analysis methods. Second, given that participants were trained to a high-performance level before entering the scanner, there was little variability in the data to correlate behavioural performance with decoding accuracy, nor did we collect data regarding training performance. This means we cannot comment on the emergence of HD representations, or how a landmark’s perceived stability correlates with fMRI measures of HD coding. Finally, participants were trained to focus on GHD meaning that our results are not sensitive to regions in which the HD activity is modulated by attentional demands.

To summarize, our multivariate reanalysis of Shine et al. (2016) provides greater insight into the neural underpinnings of coding global and local HD reference frames in the human brain. Our findings support recent rodent and human fMRI data and point to potential organisational principles of the HD signal particularly with regards to LHD.

## Acknowledgements

The authors would like to thank Dr Carl Hodgetts for helpful feedback on the manuscript.

